# A rabbit monoclonal antibody against the antiviral and cancer genomic DNA mutating enzyme APOBEC3B

**DOI:** 10.1101/513341

**Authors:** William L. Brown, Emily K. Law, Michael A. Carpenter, Prokopios P. Argyris, Rena Levin-Klein, Alison N. Ranum, Amy M. Molan, Colleen L. Forster, Brett D. Anderson, Lela Lackey, Reuben S. Harris

## Abstract

The DNA cytosine deaminase APOBEC3B (A3B) is normally an antiviral factor in the innate immune response. However, A3B has been implicated in cancer mutagenesis, particularly in solid tumors of the bladder, breast, cervix, head/neck, and lung. Here, we report data on the generation and characterization of a rabbit monoclonal antibody (mAb) for human A3B through a series of immunoblot, immunofluorescence microscopy, and immunohistochemistry experiments. Positive results indicate that one mAb, 5210-87-13, will be useful for fundamental research in virology and cancer biology and that immunohistochemical quantification of A3B enzyme levels may constitute a clinical biomarker for tumor evolvability and differential treatment.

## Introduction

The APOBEC3 family of single-stranded DNA cytosine deaminases constitutes a vital arm of the innate immune response that serves to restrict the replication of viruses and transposable elements (reviewed by refs.^1–5^). A wide range of parasites have proven susceptible to restriction and/or mutation by APOBEC3 enzymes including retroviruses and retrotransposons with obligate single-stranded DNA replication intermediates, hepadnaviruses such as HBV, small DNA tumor viruses such as human papillomavirus (HPV) and polyomaviruses (BK-PyV/JC-PyV), and large DNA viruses such as the γ-herpesviruses EBV and KSHV (*e.g*., refs.^6–16^). Thus, the human APOBEC3 enzymes have become an important focus of research in many areas of virology.

At least one human APOBEC3 enzyme has emerged recently as major endogenous source of mutation in a wide variety of different cancers (reviewed by refs.^3,5,17–20^). APOBEC mutations in cancer are identified through a hallmark signature of C-to-T and C-to-G base substitutions in 5’-TCA/T trinucleotide motifs. This APOBEC mutation signature is evident in over half of all human cancer types, and it is one of the dominant etiologies in bladder, breast, cervix, head/neck, and lung cancer^21–24^. APOBEC3B (A3B) is the leading candidate for APOBEC mutagenesis in cancer due to several independent lines of evidence including overexpression in tumors and cancer cell lines, nuclear localization, modulation (upregulation and/or counteraction) by tumor-promoting viruses including HPV, and associations with poor clinical outcomes (*e.g*., refs.^11,16,22,25–35^). APOBEC3H (A3H) may also contribute to cancer mutagenesis, particularly in populations that lack A3B due to a naturally occurring deletion allele^36,37^. Moreover, APOBEC3A (A3A) has also been implicated through a number of lines of investigation but studies on the endogenous enzyme have yet to be done in model systems due to low/no expression (*e.g*., refs.^23,38–46^). Ongoing research by many groups is likely to clarify the relative contributions of each APOBEC enzyme to mutagenesis in different tumor types, and new technologies and reagents including validated monoclonal antibodies (mAbs) are likely to have essential roles in this process.

A problem studying human APOBEC3 enzymes in virology and cancer biology is the fact that human cells have the potential to express up to seven different family members, A3A, A3B, and A3H, as described above, as well as APOBEC3C (A3C), APOBEC3D (A3D), APOBEC3F (A3F), and APOBEC3G (A3G). Moreover, due to multiple primate-specific gene duplication and diversification events, several of the present-day human A3 enzymes share high levels of amino acid identity^47,48^. For instance, A3A and the catalytic domain of A3B are over 90% identical, and each of these proteins shares over 65% identity with the catalytic domain of A3G. Thus, it has also been challenging to develop mAbs to distinguish between these enzymes in various immunoassays.

Here we report the development of a rabbit mAb called 5210-87-13 for human A3B. This reagent has proven effective in a variety of different immunoassays including enzyme-linked immunosorbent assay (ELISA), immunoblot (IB), immunofluorescent microscopy (IF), flow cytometry (FLOW), and immunohistochemistry (IHC). However, despite clear demonstrations of highly significant and broad utility, a remaining liability for consideration when interpreting 5210-87-13 data is potential for cross-reactivity against A3A and A3G, which stems from unavoidable homology in the original peptide used for immunization.

## Results

### Epitope design and hybridoma generation

The full protein sequence of the seven human APOBEC3 enzymes was obtained from GenBank. ClustalW was used to identify regions unique to A3B, and a C-terminal region was selected for synthesis of a peptide immunogen corresponding to residues 354-382. Although this 28 residue sequence is unique to A3B, homology with related family members was unavoidable and the homologous regions of A3A and A3B still shared 27/28 and 25/28 residues, respectively (Fig. 1A). The selection of this peptide was also encouraged by knowledge that the corresponding C-terminal peptide of A3G is immunogenic in rabbits^49,50^.

**Figure 1.**
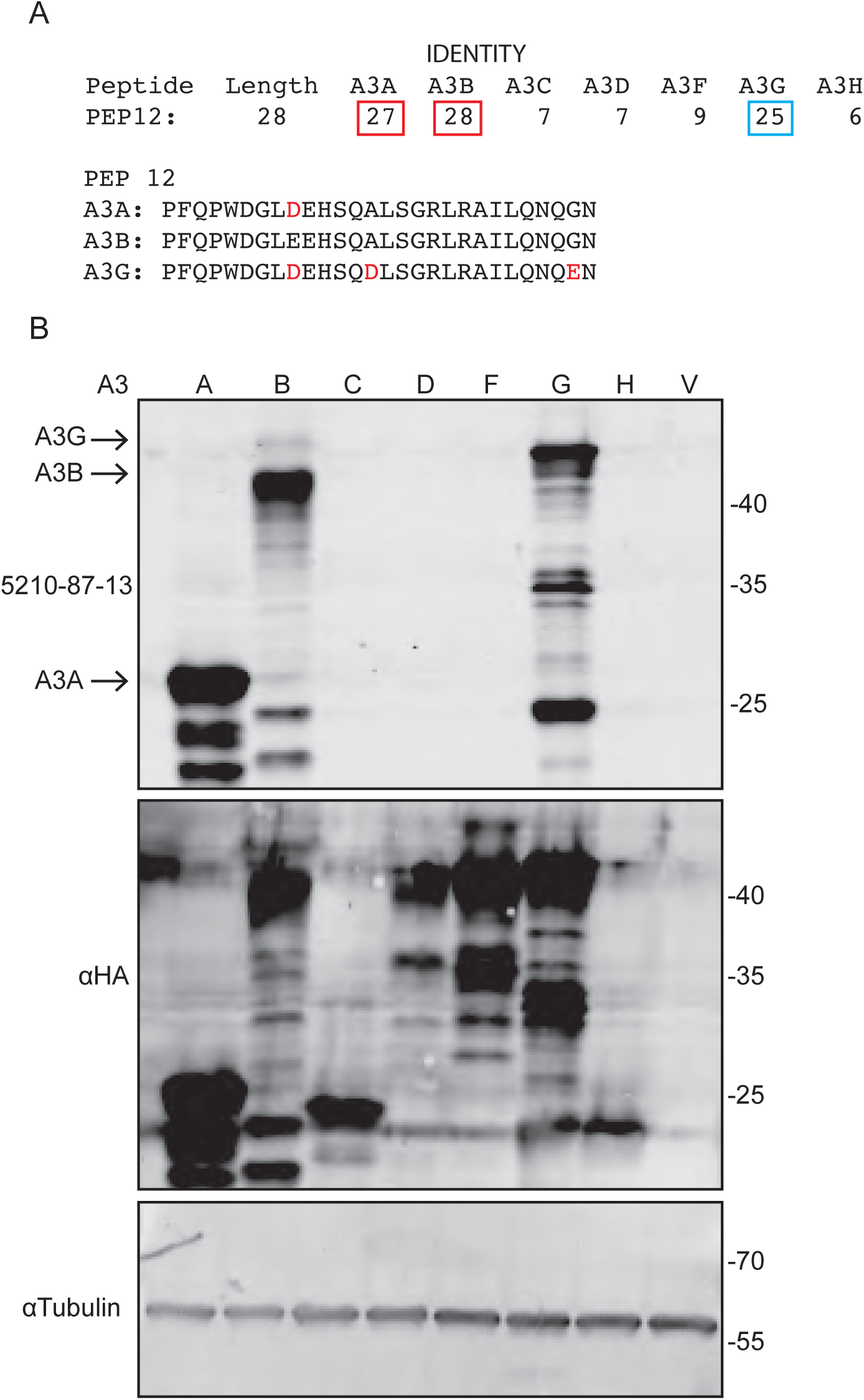
Peptide selection and anti-A3 reactivity of the 5210-87-13 mAb by immunoblot. **A.** A3B C-terminal peptide (PEP12) identity across human A3 family members. **B.** Anti-A3 reactivity of 5210-87-13, with anti-HA and anti-Tubulin blots as expression and loading controls, respectively.

The A3B peptide 354-382 was used to immunize two rabbits over a period of 10-12 weeks. The initial immunization occurred through 3 injections of KLH-conjugated peptide, and an additional boost was done through 2 injections of OVA-conjugated peptide. Throughout the immunization protocol, sera were assessed for anti-A3B activity by ELISA, immunoblot, and immunofluorescence microscopy assays (data not shown). Both rabbits (5210, 5211) showed positive anti-A3B immune responses and were given a final antigen boost before spleens were harvested for B cell isolation and hybridoma production.

Splenic B lymphocytes were fused with the rabbit plasmacytoma cell line 240E-W to create hybridoma candidates (40 x 96-well plates). Standard ELISA assays were used to screen culture supernatants from 3840 candidate wells for immunoreactivity against recombinant A3B C-terminal domain^51^. Positive candidates were expanded and supernatants were retested by ELISA and further screened by IB and IF (data not shown). Finally, the strongest ELISA-positive A3B-reactive candidates were subcloned by limiting dilution to create monoclonal hybridoma cell lines for further characterization.

Sera from expanded monoclonal hybridomas were next tested for specificity against human A3B by immunoblotting. A3B-low 293T cells^16^ were transiently transfected with HA-tagged A3 expression constructs (A3A-HA, A3B-HA, A3C-HA, A3D-HA, A3F-HA, A3G-HA) or the empty expression vector alone as a negative control. After a 24 hr incubation for protein expression, whole cell extracts were prepared, fractionated by SDS-PAGE, and probed using supernatants from each candidate hybridoma. Several ELISA-positive and IB-positive mAb candidates were obtained and all showed varying degrees of reactivity against A3B, as expected, but also varying degrees of reactivity against A3A and A3G. The hybridoma clone 5210-87-13, which produced the strongest A3B-reactive mAb was selected for further characterization (upper panel, Fig. 1B). Control anti-HA immunoblots confirmed expression of each A3-HA construct, and control anti-Tubulin immunoblots showed similar protein loading in each lane (lower panels, Fig. 1B).

### Immunofluorescent microscopy results using the 5210-87-13 anti-A3B mAb

Prior studies have shown that the bulk of cellular A3B is nuclear due to an active import mechanism (*e.g*., refs.^25,52–54^). To determine whether the 5210-87-13 anti-A3B mAb recognizes nuclear-localized A3B, HeLa cells were transfected with an A3B-eGFP expression construct and subjected to IF microscopy. A3B-eGFP-expressing HeLa cells showed strong green fluorescent signal in the nucleus (Fig. 2A). In comparison, staining the same cells with the 5210-87-13 mAb followed by a secondary goat anti-rabbit-IgG-TRITC resulted in red signal from the same nuclear regions (Fig. 2A). Signal co-localization was confirmed in the merged image with staining of the entire nucleus except nucleoli. Similar results were obtained with T-REx-293-A3B-eGFP cells treated with doxycycline to induce A3B-eGFP expression (Fig. 2B), except a broader cell-wide co-localization was observed likely due to the combined effects of gross overexpression and a previously reported nuclear import defect in 293 and derivative cell lines^52,55^.

**Figure 2.**
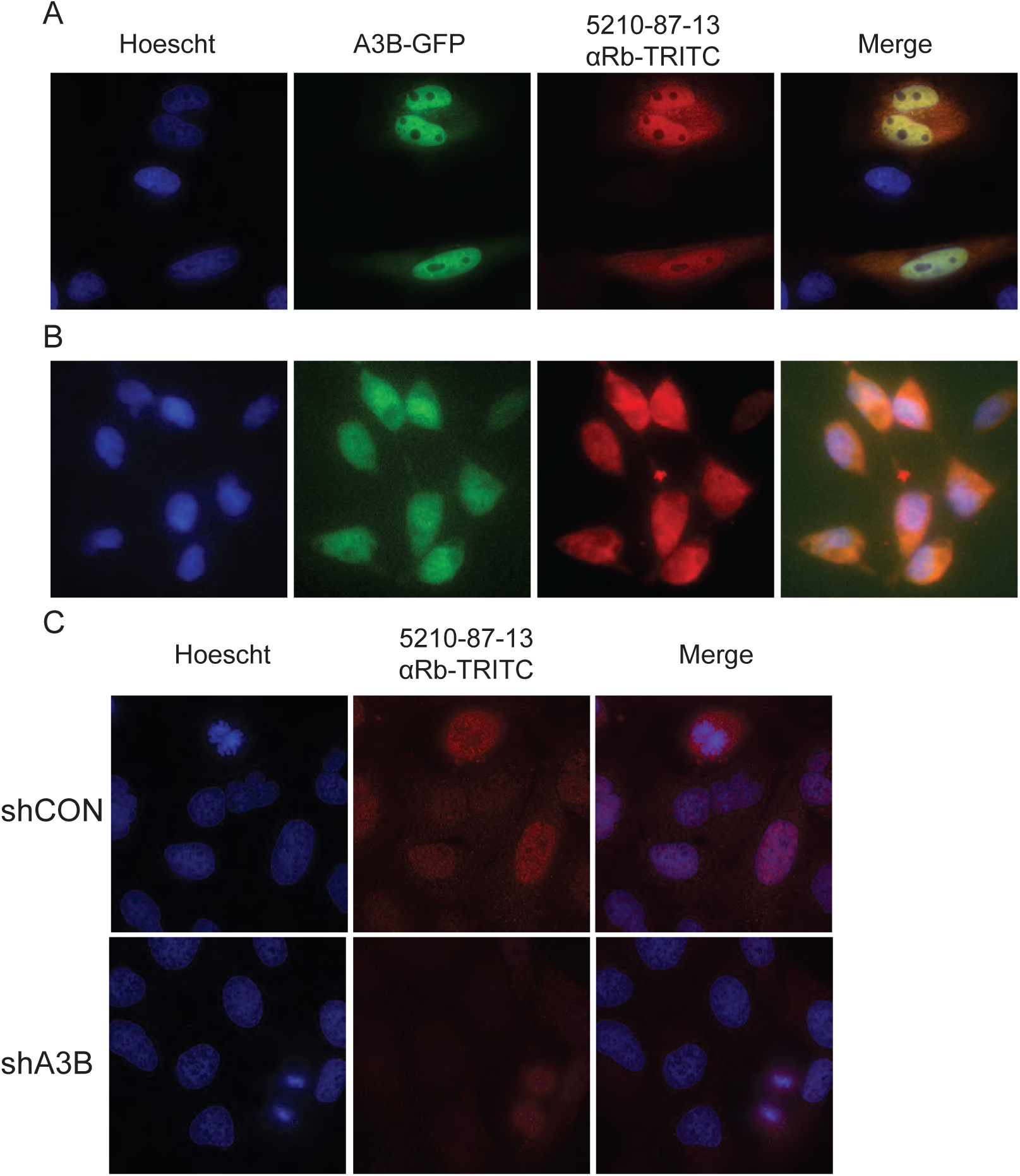
Immunofluorescent microscopy results for the 5210-87-13 mAb. **A.** IF microscopy images of HeLa expressing A3B-eGFP and stained with the 5210-87-13 mAb. **B.** IF microscopy images of doxycycline-induced T-REx-293-A3B-eGFP cells stained with the 5210-87-13 mAb. **C.** IF microscopy images 5210-87-13 stained U2OS cells expressing a control shRNA (shCON) or shA3B.

Next, immunofluorescent microscopy was used to ask whether the 5210-87-13 anti-A3B mAb could detect endogenous A3B. U2OS osteosarcoma cells were transduced with shRNA against *A3B* to knockdown endogenous *A3B* gene expression or with scrambled shRNA to generate a control cell population, respectively. A3B-depleted and control U2OS cells were then stained as above with 5210-87-13 mAb (primary) and anti-rabbit-IgG-TRITC (secondary) and subjected to fluorescent microscopy. A3B-depleted cells were negative or stained poorly, while the control population with endogenous A3B showed red nuclear fluorescent signal largely coincident with Hoescht (Fig. 2C). As previously reported, A3B is largely cell wide and excluded from chromatin during mitosis^52,53^ (*e.g*., top cell in shCON image in Fig. 2C). Taken together, these immunofluorescent microscopy data show that the 5210-87-13 anti-A3B mAb is capable of labeling both transiently and endogenously expressed A3B in different cell types.

### Immunoblot results using cancer cell lines and the 5210-87-13 anti-A3B mAb

Given strong A3B overexpression in various tumor types (see Introduction), a small panel of cancer cell lines was selected for additional 5210-87-13 anti-A3B IB validation experiments. First, two multiple myeloma cell lines (NCI-H929 and JJN3) and two primary effusion lymphoma-derived cell lines (JSC-1 and BC-3) were tested due to RNAseq data indicating relatively high *A3B* expression^56^. These immunoblots showed strong reactivity around 36 kDa relative to the molecular weight size marker, which is the position where endogenous A3B is routinely observed migrate (Fig. 3A). Endogenous A3G migrates slightly slower with, for instance, BC-3 expressing high A3B and low A3G and the other lines expressing similar levels of both proteins. The identity of the 36 kDa anti-A3B band was confirmed in JJN3 cells through CRISPR knockout of the endogenous *A3B* gene (Fig. 1A). It is important to note that both A3B and A3G migrate faster in SDS-PAGE gels than their predicted molecular weights (45 and 46 kDa, respectively) and that separation can be facilitated by gel type and run duration (see Methods).

**Figure 3.**
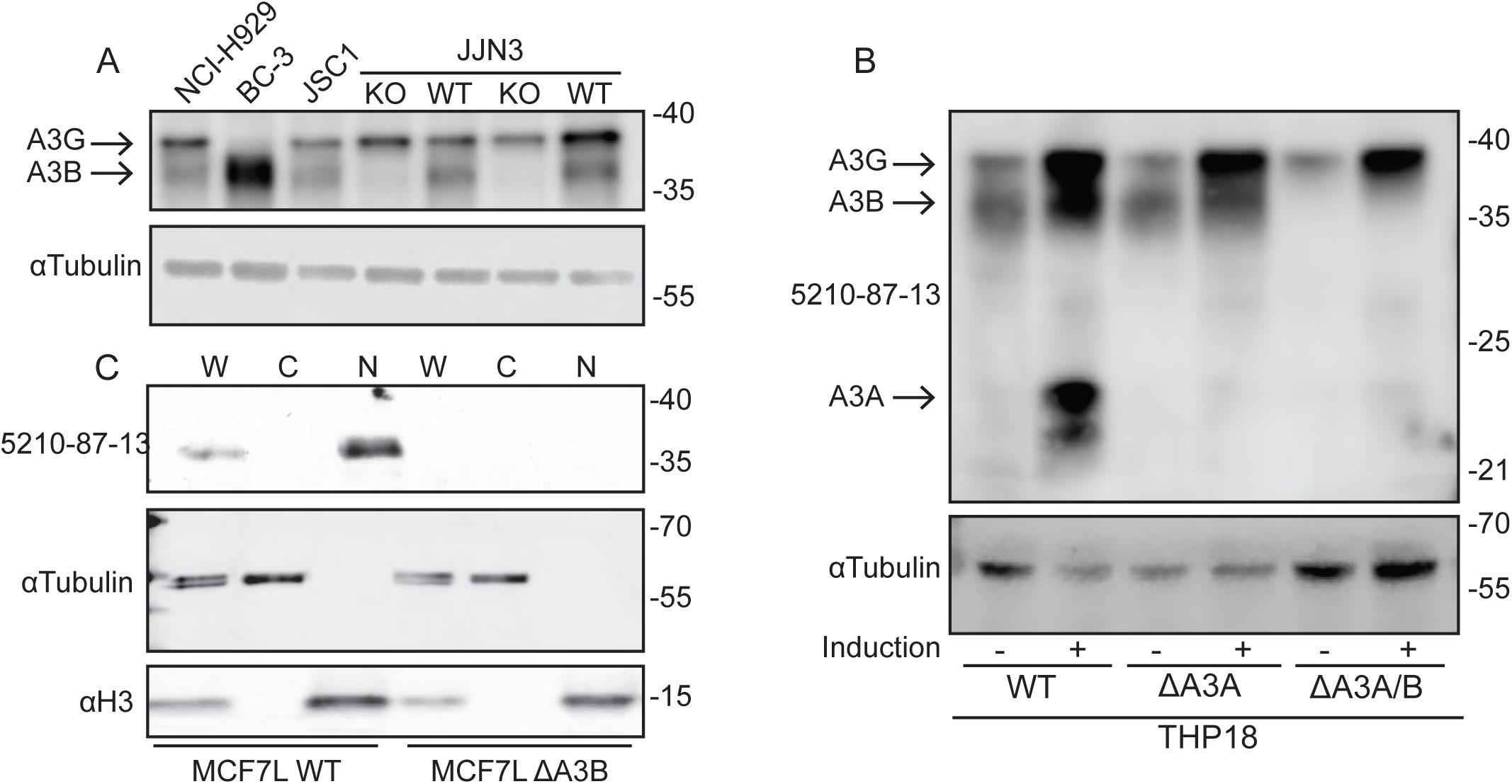
Endogenous A3B detection by 5210-87-13 mAb immunoblotting. **A, B.** Immunoblots of whole cell lysates from the indicated cell lines probed with the 5210-87-13 mAb and anti-Tubulin as a loading control. **A, B.** Immunoblots of whole cell lysate, cytoplasmic extract, and nuclear extract from the indicated cell lines probed with the 5210-87-13 mAb and anti-Tubulin as a loading control.

Additional validation studies were done using THP18, a subclone of the monocytic cell line THP1, and derivative lines engineered by CRISPR to lack *A3A* or both *A3A* and *A3B*. THP1 is known to express A3B and A3G in normal growth medium and, following interferon-α (IFN-α) treatment, induced levels of A3A^50,57,58^. These lines were expanded in normal growth medium, treated with IFN-α (330 u/mL) and LPS (1 μg/ml), processed into whole cell extracts, and analyzed by immunoblotting with the 5210-87-13 mAb (Fig. 3B). The resulting immunoblot shows that A3A, A3B, and A3G are all induced under these conditions, with A3A undergoing the most dramatic change from undetectable to a robust visible signal (2 bands due to alternative methionine codons for translation initiation, as described^50,57,58^). Moreover, A3A expression was abolished in the *A3A* knockout, and both A3A and A3B expression were abolished in the double knockout (THP18 *∆A3A/B*, Fig. 3B).

Last, nuclear and cytoplasmic fractions from a previously described MCF-7L breast adenocarcinoma cell line and a derivative engineered by CRISPR to lack A3B^36^ were analyzed by immunoblotting with the 5210-87-13 mAb (Fig. 3C). This experiment showed as expected that A3B is detectable in whole cell extracts, absent from cytoplasmic fractions, and enriched in nuclear lysates and, importantly, completely undetectable in engineered *A3B* knockout cells. In comparison, tubulin is enriched in cytoplasmic fractions and histone H3 in nuclear fractions. No cross-reactivity with A3G or A3A was observed in MCF-7L because this it does not express significant levels of the mRNAs that encode these proteins under normal growth conditions^36^.

### Immunohistochemical detection of A3B in cell lines using the 5210-87-13 anti-A3B mAb

We next wanted to evaluate the efficacy of the 5210-87-13 anti-A3B mAb in IHC experiments using a custom cell microarray (CMA). As described above, we have both identified and purposefully engineered systems to express the broadest possible range of A3B protein from absolute zero (*A3B* knockout cell lines) to massive overexpression (T-REx-293-A3B-eGFP cells treated with doxycycline). Two isogenic sets of cell lines were selected for these IHC experiments: 1) 293T and T-REx-293-A3B-eGFP minus/plus doxycycline induction and 2) THP18, THP18 *∆A3A* and THP18 *∆A3A/B* minus/plus IFN-α/LPS induction. In all instances, cell lines were grown under normal conditions, treated to induce *A3B* or *A3A/B/G* expression, and then split into aliquots for mRNA quantification by RTqPCR using established assays^25,36,57,59^ and for protein detection by IHC using the 5210-87-13 mAb. The RTqPCR results largely mirrored the immunoblot data in Fig. 1A for 293T, described previously for the T-REx-293-A3B-eGFP system^25,58,60^, and shown above in Fig 3B for the THP18 cell line and its *A3A*- null and *A3A/B*-null derivatives. *A3A* mRNA is undetectable in the 293T cells (regardless of treatment) and induced by 3 orders of magnitude in IFN-α/LPS treated THP18 and, of course, absent from *A3A*-null and *A3A/B*-null derivatives (Fig. 4A). *A3B* mRNA is weakly expressed in 293T and modestly expressed in untreated T-REx-293-A3B-eGFP cells (due to leaky expression); in contrast, doxycycline treatment induced *A3B* mRNA levels by 2 orders of magnitude (Fig. 4A). *A3B* mRNA levels decreased modestly following IFN-α/LPS treatment despite clear increases in protein levels in IB experiments, further emphasizing the need for protein-level quantification (compare data in Fig. 4A and Fig. 3B). In comparison, *A3G* mRNA is expressed weakly in the 293T lines (undetectable by IB) and induced by IFN-α/LPS treatment in THP18 and most derivatives, consistent with IB results (Fig. 4A and Fig. 3B).

**Figure 4.**
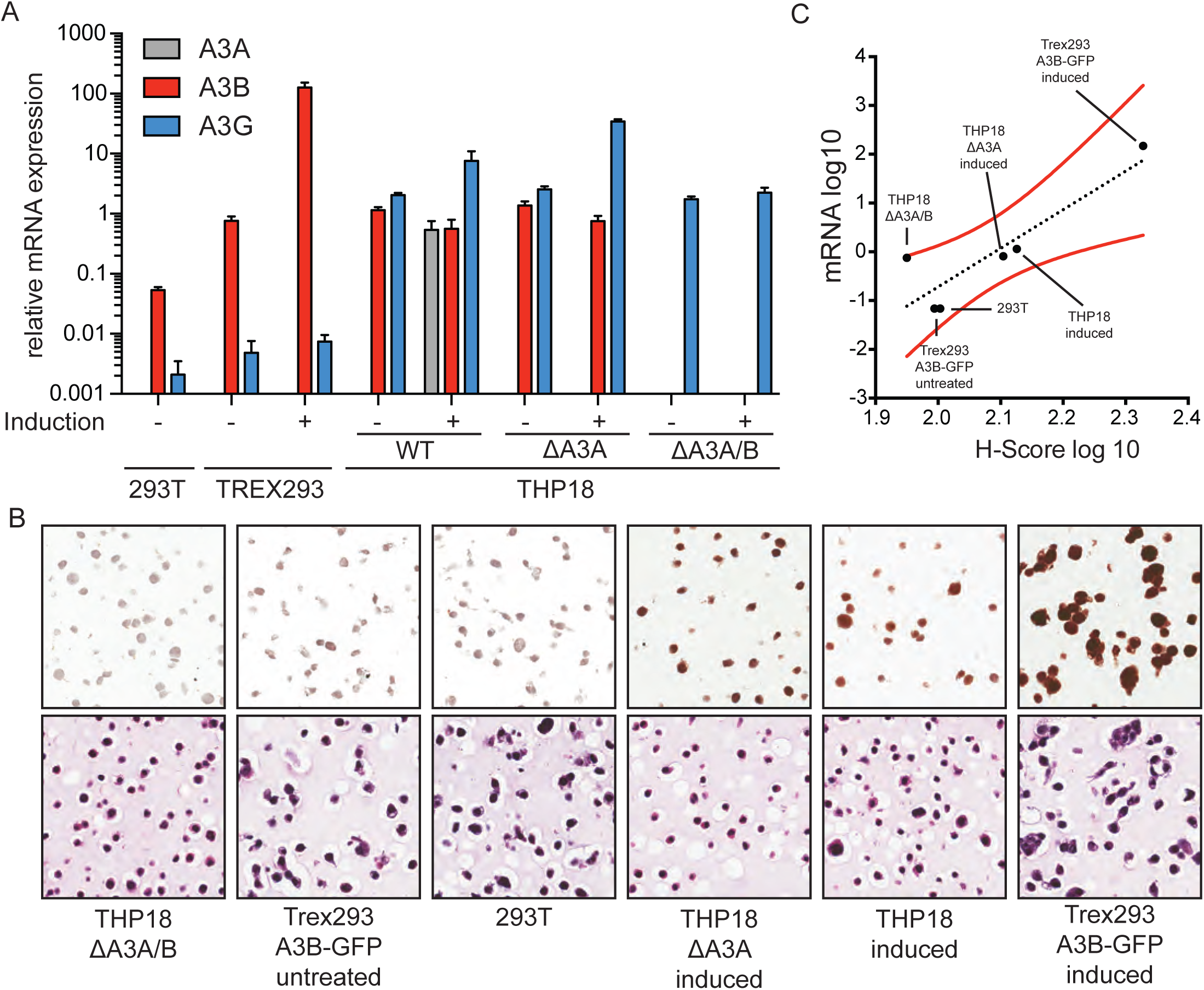
*A3B* mRNA levels correlate with IHC H-score using the 5210-87-13 mAb. **A.** Assessment of *A3A*, *A3B*, and *A3G* mRNA levels by RTqPCR in the indicated cell lines. **B.** Representative IHC images from 5210-87-13 staining of the indicated cell lines in the context of a CMA (top row) along with corresponding hematoxylin and eosin (H&E) photomicrographs. **C.** A dot plot showing the positive correlation between *A3B* mRNA levels and IHC H-score from 5210-87-13 mAb nuclear straining of the indicated cell lines.

In IHC experiments, doxycycline-induced T-REx-293-A3B-eGFP cells showed the strongest nuclear A3B immunoreactivity in almost 100% of the examined cells (far right images in Fig. 4B). On the other end of the signal intensity scale, the *A3A/B*-null THP18 cell line exhibited sporadic and weak immunostaining with the 5210-87-13 mAb, suggesting that the potentially cross-reactive epitope in A3G may be either blocked or shielded under these experimental conditions (far left images in Fig. 4B). Consistent with this interpretation, IFN-α/LPS treated THP18 and *A3A*-null derivatives exhibited the 2^nd^ and 3^rd^ highest (strong to moderate) nuclear A3B immunoreactivities, respectively. In comparison, untreated parental 293T and T-REx-293-A3B-eGFP cells exhibited sporadic and weak immunostaining. Despite likely cross-reactivity with A3A and little/no cross reactivity under these conditions with A3G, a strong positive correlation was evident between *A3B* mRNA levels and 5210-87-13 IHC Histo-score (H-score; R^2^ = 0.8, Pearson correlation coefficient=0.9; Fig. 4C).

### Immunohistochemical detection of endogenous A3B in tumor tissue using the 5210-87-13 anti-A3B mAb

Last, we sought to determine whether the 5210-87-13 mAb can detect A3B protein in formalin-fixed and paraffin embedded (FFPE) patient biopsy specimens. Tumor tissues were chosen to reflect cancers in which *A3B* mRNA overexpression has been reported, specifically cancers of the cervix, breast, bladder, and head/neck^25^. In many instances, the 5210-87-13 mAb specifically and clearly stained the nuclear compartment of tumor cells but not surrounding non-tumor cells (Fig. 5A and data not shown). However, in tumor cases where positive nuclear staining was evident, considerable intra-tumor heterogeneity was observed with respect to both distribution and intensity (*e.g*., a strong A3B-positive tumor cell may be next to an A3B-negative tumor cell). A molecular explanation for this heterogeneity is not available at this time but the observation again underscores the need for protein-level detection methods for A3B in cancer.

**Figure 5.**
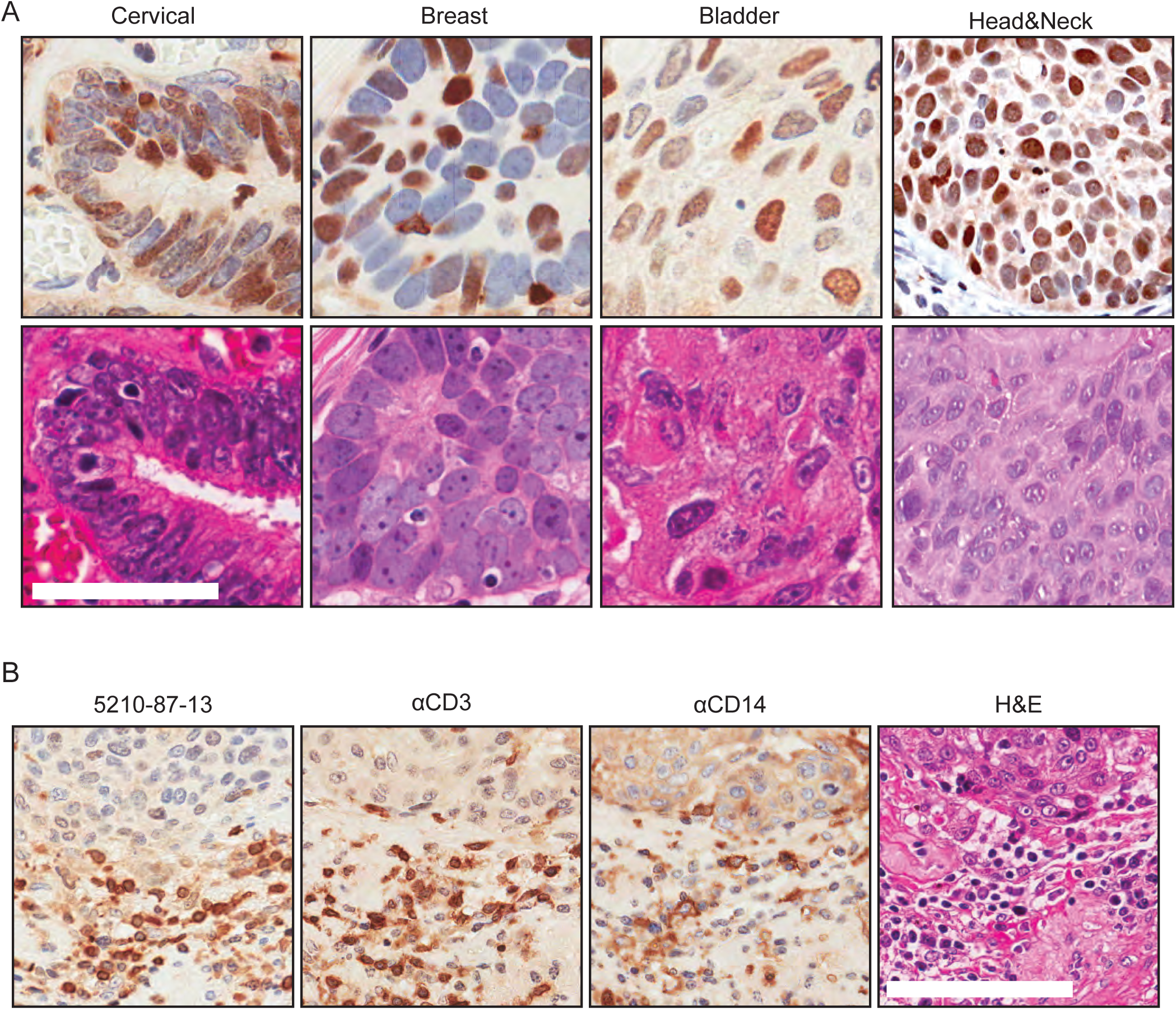
IHC detection of endogenous A3B protein in tumors using the 5210-87-13 mAb. **A.** 5210-87-13 mAb staining of the indicated FFPE tumor specimens prescreened by RTqPCR for high *A3B* mRNA levels (top row) with serial H&E-stained sections (bottom row). **B.** 5210-87-13 mAb, anti-CD3, and anti-CD14 staining of a representative FFPE tumor specimen with inflammation and infiltrating immune cells. See text for details.

In addition, in a subset of tumors showing evidence of inflammation, the surrounding connective tissue stroma was infiltrated by a mixed population of CD3-positive and CD14-positive inflammatory cells (Fig. 5B). CD3 and CD14 are diagnostic markers for T-lymphocyte and monocytic lineage cells, respectively. Interestingly, in these cell types the strongest 5210-87-13 mAb reactivity was noticed in the cytoplasmic compartment, where A3G and A3A are known to be localized ^7,58,61–63^. Thus, these observations combined indicate that 5210-87-13 mAb has the capacity to label nuclear A3B in neoplastic cells, and it most likely reacts with cytoplasmic A3G and A3A in tumor infiltrating lymphocytes (TILs) and tumor-associated monocytes/macrophages, respectively.

## Discussion

Herein, we report the creation and validation of a rabbit anti-human A3B mAb (5210-87-13). This mAb is active in a wide variety of immunoassays including ELISA, IB, IF, and IHC. As expected from positive IF results, it is also useful for FLOW (data not shown). Despite our original goal of identifying a anti-A3B-specific mAb, the 5210-87-13 mAb and all other hybridoma candidates still cross-reacted to varying degrees with A3A and A3G. However, data from experiments utilizing the 5210-87-13 mAb are still interpretable, and in many instances definitive, provided 1) A3B is the only one of these enzymes expressed in the system (*e.g*., many different cancer cell lines and tumors), 2) the three enzymes can be distinguished by kilodalton size and mobility (immunoblots), or 3) the enzymes localize to different cell types or subcellular compartments (A3B shows nuclear localization whereas A3A and A3G are cytoplasmic). The experiments reported here further demonstrate that cross-reactivity with more distantly related human APOBEC3 family members will not be a complicating issue.

The 5210-87-13 mAb detailed here has already proven effective in a range of IB experiments. We first used it to study induction of endogenous A3B expression by the PKC/non-canonical NF-κB signal transduction pathway^27^. We subsequently used it in IB experiments to study SIV Vif-mediated degradation of endogenous A3B^64^, endogenous A3B upregulation by polyomavirus infection^11^, A3B lysine post-translational modification^65^, and endogenous A3B interaction with EBV BORF2^16^. The 5210-87-13 mAb has also been shared with colleagues, who have independently demonstrated utility in anti-A3B IB experiments^66,67^. In addition, another independent study used 5210-87-13 to monitor A3A and A3B induction in IB and IF experiments^68^. Thus, depending on the experimental system and biological question, the 5210-87-13 mAb detailed here may be useful for studying human A3A, A3B, and A3G.

Equally important, the studies here are the first to use the 5210-87-13 mAb to measure A3B protein in FFPE-tumor tissues using IHC. This technique opens the door to a wide range of clinical studies using 5210-87-13 to assess the diagnostic and/or prognostic utility of nuclear A3B levels in multiple human malignancies, including bladder, breast, cervix, head/neck, and lung cancer. For instance, because high-risk HPV infection can cause A3B upregulation^28,29,69^, nuclear IHC staining using 5210-87-13 may be useful as a sentinel marker (or surrogate marker in addition to p16) for diagnosing HPV-related precancerous or cancerous lesions in infected cervical, head/neck, and bladder tissue. In addition, consistent with initial results in breast cancer using mRNA expression levels^30,33,70^, tumors immunoreactive for A3B using 5210-87-13 may have higher degrees of evolvability in comparison to those that are A3B-negative and exhibit increased rates of metastasis and drug resistance.

## Materials and Methods

### Cell lines

293T, HeLa, and U2OS were cultured in RPMI supplemented with 10% fetal bovine serum (FBS, Thermo Fisher Scientific) and 1% penicillin-streptomycin (P/S, Thermo Fisher Scientific). T-REx-293T is a derivative of 293T engineered to constitutively expresses the tetracycline repressor^25,58,60^. THP18^71^ was derived from a single cell subclone of the monocyte cell line THP-1 (ATCC TIB-202), and maintained in the same growth medium as above. MCF-7L and its A3B-null derivative have been described^33,36^, and these lines were maintained in IMEM supplemented with 15% FBS and 1% P/S, non-essential amino acids (0.1 mM), and insulin (11.25 nM). Multiple myeloma cell lines NCI-H929 and JJN3 were kind gifts from Brian Van Ness (University of Minnesota Twin Cities) and were grown in RPMI-1640 supplemented with 15% FBS, 1% P/S and 50 µM β-mercaptoethanol. PEL cell lines JSC-1 and BC-3 were kind gifts of Bill Sugden (University of Wisconsin Madison) and grown in RPMI-1640 supplemented with 10% FBS and 1% P/S.

### Plasmids

The A3-HA plasmid set has been described^7^. The A3B-eGFP expression construct, and the shRNA expression constructs have also been reported^25,52,53^.

### *Generation of* A3A *and* A3A/B *knockout cell lines*

*A3A* and *A3A/B* deletion THP18 cell lines and *A3B* deletion JJN3 cells were engineered by co-transduction with pLentiCRISPR expressing the gRNA sequences flanking the *A3A* gene (CCTGGACAAGCGACATACCG and AATGCACCATGTCTCCCCTC), both the *A3A* and *A3B* genes (CCTGGACAAGCGACATACCG and ACAACACCCTCGCCCCATGA), or the *A3B* gene alone (AATGCACCATGTCTCCCCTC and ACAACACCCTCGCCCCATGA). As control, the parental THP18 cell line was transduced in parallel with an unrelated gRNA (GGCGACCACCGCCGCCATCT). 3 days following transduction, cells were treated with 1 µg/mL puromycin for 3 days, and seeded for single cell cloning in 96 well plates 1 week later. Deletion mutant lines were identified by PCR using primers amplifying within the *A3B* gene ^72^ (TTGGTGCTGCCCCCTC and TAGAGACTGAGGCCCAT) and the *A3A* gene (CCTCCTCTGGTCTTTTCCCT and GAAACCACAAGTACAATCCGG) and confirmed by qPCR and immunoblots.

### Hybridoma generation

The full protein sequences of the seven human APOBEC3 (A3) enzymes were obtained from GenBank (A3A GenBank: AAI26417.1; A3B GenBank: AAW31743.1; A3C GenBank: AAH11739.1; A3D GenBank: AIC57731.1; A3F GenBank: AAZ38720.1; A3G GenBank: AAZ38722.1; A3H GenBank: ACK77774.1). ClustalW was used to identify regions unique to A3B. Residues 354-382, PFQPWDGLEEHSQALSGRLRAILQNQGN, were used to create a peptide immunogen (Epitomics).

Two rabbits were given 3 injections using KLH-conjugated peptide then 2 further injections with OVA-conjugated peptide over the course of 10-12 weeks (Epitomics). Test bleeds from the rabbits were screened at UMN for anti-A3B expression by immunoblot with lysates from 293T cells that expressed A3 proteins tagged with the HA epitope. The bleeds were further screened at UMN by immunofluorescence microscopy (IF) of HeLa cells expressing A3B-eGFP protein. Rabbits showing positive anti-A3B immune responses were selected for a final immunization boost before the spleens were harvested for B cell isolation and hybridoma production. Hybridoma fusions of 240E-W cells with lymphocytes from the selected rabbits were performed by Epitomics in 40 x 96-well plates. Cell supernatants were screened at UMN by A3B ELISA and the strongest positive hybridoma pools were subcloned by standard limiting dilution to generate monoclonal hybridoma cell lines. Hybridoma 5210-87-13 was expanded to 1L and then switched to a serum-free medium for one week. This medium was clarified by centrifugation to remove cells and then passed over a Protein A column to bind mAb. The resulting mAb was eluted in glycine pH 2.5 and dialyzed into PBS.

### ELISA

Standard ELISA screening with recombinant A3Bctd^51^ was used to monitor anti-peptide immune responses in rabbits and to identify single-clone hybridomas that expressed anti-A3B antibodies. Recombinant A3Bctd (20 ng/well) was immobilized on 96-well ELISA plates, blocked with 3% BSA, then incubated with undiluted cell-free media supernatant from candidate hybridomas. Supernatants from cells that did not express A3B binding activity were used as negative controls. The positive control was a rabbit anti-human A3G antibody (NIH AIDS Reagent Program, 10201) that cross-reacts with A3B. Binding was detected with a goat anti-rabbit HRP secondary antibody (Jackson Immunodiagnostics1:5000), visualized with tetramethylbenzidine (TMB), and quantified by spectroscopy at 450 nm.

### Immunoblots

Cell lysates from 293T cells, transiently transfected with each HA-tagged A3 protein or the expression vector alone, were resolved by 12% SDS-PAGE and transferred to PVDF membrane. Membranes were probed with the test bleeds (1:1000), cell-free supernatants from each hybridoma cell line (1:3), anti-HA (C29F4, #3724, Cell Signaling Technology, Danvers, MA, 1:1000), or anti-tubulin (MMS-407R, Covance, Emeryville, CA, 1:20,000) in 50% BLOK (Millipore), 0.1% Tween 20 in PBS. The membranes were then incubated with secondary antibody, either goat anti-rabbit-HRP (Jackson 1:5000), anti-rabbit IgG IR800CW (Odyssey 926-32211, 1:20,000), or anti-mouse IgG IR800CW (Odyssey 827-08364, 1:20,000) in 50% BLOK (Millipore), 0.1% Tween 20 in PBS. Signals were detected with HyGlo (Denville Scientific) on film, or imaged using LiCor.

### Immunofluorescence microscopy

HeLa cells (2×10^5^) were transfected with 200 ng an A3B-eGFP expression construct. T-REx-293-A3B-eGFP cells (2×10^5^) were induced with 1 μg/mL doxycycline. After 24 hrs, cells were washed twice with PBS, fixed in 4% paraformaldehyde (PFA) for 30 min, washed twice with PBS, then incubated overnight at room temperature with primary antibody in blocking buffer (5% goat serum, 1% BSA, and 0.2% TritonX-100). Supernatant from 5210-87-13 hybridoma cell culture was used at 1:5 dilution, and purified anti-A3B (5210-87-13) at 1:250 dilution in blocking buffer. Cells were washed twice with 1x PBS and incubated in secondary antibody, anti-rabbit-TRITC (1:500 Jackson ImmunoResearch Laboratories Inc. 111095144), or anti-rabbit-Alexafluor 594 1:500 in blocking buffer for 1 hr at room temperature. Following two washes with PBS, the nuclei were strained with 0.1% Hoescht for 15 minutes at room temperature. Following two washes with 1x PBS, slides were mounted in 50% glycerol. Images were taken at 60x magnification, with a 1 second TRITC exposure time (to normalize), cropped to 900×900 pixels, and scaled down to 10% size.

### Immunohistochemistry

Human tissues were obtained using a protocol that was approved by the University of Minnesota Institutional Review Board. All formalin-fixed, paraffin-embedded (FFPE) specimens were derived from incisionally or excisionally biopsied cervical, breast, bladder, and head and neck carcinomas. Tissues were sectioned at 4 µm, mounted on positively charged, adhesive slides and allowed to air-dry for at least 24 hrs. To deparaffinize and rehydrate the samples, slides were baked in a 65°C oven for 20 min, washed 3 times with CitrisolvTM (Decon Labs, #1601) or xylene for 5-min/each, soaked in graded alcohols (100% x 2, 95% and 80% for 3 min/each), and then rinsed in running water for at least 5 min. Epitope retrieval was performed using Reveal Decloaker (BioCare Medical, # RV1000M) in a steamer for 35 min, followed by a 20 min “cool-down” period. Then, slides were rinsed with running tap water for 5 min and transferred to TBST for 5 min. Endogenous peroxidase activity was quenched by placing the slides in 3% H_2_O_2_ in TBST for 10 min at RT, followed by a 5-min rinse under running water. To block non-specific binding of primary antibody, sections were covered with Background Sniper (BioCare Medical, # BS966MM) for 15 min at RT. After blocking, serial sections of each tumor were incubated overnight at 4°C with 5210-87-13 mAb diluted (1:300-1:1,000) in 10% Sniper in TBST. Immunostaining against cells of lymphocytic and monocytic lineage was performed using anti-CD3 and anti-CD14 antibodies, respectively.

Following overnight incubation with primary antibody, sections were rinsed in TBST for 5 min, and completely covered with post-primary rabbit anti-mouse IgG (Leica Biosystems, Novolink, # RE7260-K) for 30 min at RT. Then, slides were drip dried, transferred to TBST for 5 min and incubated with anti-rabbit poly-HRP-IgG (Leica Biosystems, Novolink Polymer, #RE7260-K) for 30 min at RT. The reaction product was developed using the Novolink DAB substrate kit (Leica Biosystems, # RE7230-K) at RT for 5 min, rinsed in tap water for 5 min, counterstained in Mayer’s hematoxylin solution (Electron Microscopy Sciences, # 26252-01) for up to 5 min, dehydrated in graded alcohols and CitrisolvTM, and cover-slipped using Permount mounting media. The immunohistochemical stains were scanned at 40x magnification and visualized using the Aperio ScanScope XT system (Leica Biosystems).

### Cell microarray (CMA) and anti-A3B histoscore determination

1.0 x 10^6^ THP18 (WT), THP18-ΔA3A and THP18-ΔA3A/B cells were seeded in 6-well plates, allowed to grow for 24 hrs, and were then induced with 1 µg/ml LPS and 300 U/ml of IFN-α for 24 hrs. 293T and inducible T-REx-293-A3B-eGFP cells were also included in the CMA with the latter induced for 24 hrs with 1µg/µl doxycycline. Cell buttons were prepared as follows: Trypsinization, centrifugation, and supernatant removals were followed by 3 washes with PBS. Following the last wash, cells were resuspended in 10% formalin for 15 min. After fixation, cells were rinsed 3x with PBS, mixed with 2% agarose (Sigma, #A6013) at 65°C and allowed to solidify at RT for approximately 10 min. The resulting cell blocks were placed in Histosette II cassettes, dehydrated, and embedded in paraffin.

Immunohistochemical staining of the CMA was performed as above for tissues. Nuclear A3B immunoreactivity was visualized with the Aperio ScanScope XT (Leica Biosystems) and quantified using the Aperio Nuclear Algorithm. Histoscore (H-Score) was calculated using the formula (3+)×3+(2+)×2+(1+)×1 as described^73,74^.

## Acknowledgements

We thank CTSI BioNet staff including Cole Drifka and Lori Holm for tissue procurement, preparation, and sectioning, Brian Dunnett for sharing expertise regarding the use of the Aperio ScanScope XT platform and analysis of immunohistochemical staining, Matt Jarvis for assistance selecting cancer cell lines, and Bill Sugden, Brian Van Ness, and Douglas Yee for providing cell lines.

